# Gaussian accelerated Molecular Dynamics – Thermodynamic Integration (GaMD-TI): Improved alchemical free energy calculations with enhanced sampling

**DOI:** 10.64898/2026.09.01.748610

**Authors:** Yinglong Miao, Skanda Sastry, Jinan Wang, Aamir Mehmood, Michael Tae-jong Kim

**Affiliations:** Department of Pharmacology, Computational Medicine Program and Lineberger Comprehensive Cancer Center, University of North Carolina−Chapel Hill, Chapel Hill, North Carolina 27599, United States; Analytical Development and Quality Control, Genentech Inc, South San Francisco, California 94080, United States; Lingang Laboratory, Shanghai, 200031, China

**Author notes:** Corresponding authors: Y.M. and M.T.K.

**Keywords:** Free energy, Thermodynamic Integration, Gaussian accelerated Molecular Dynamics, enhanced sampling, mutation

## Abstract

It is valuable to calculate alchemical free energy changes in drug discovery and development. Thermodynamics Integration (TI) has been widely used in computational chemistry for estimating free energy changes with alchemical transformations. However, TI based on usually short Molecular Dynamics (MD) simulations often suffers from insufficient conformational sampling. Here, we have integrated Gaussian accelerated MD and TI (GaMD-TI) to enhance the conformational sampling and improve accuracy of free energy calculations. GaMD-TI has been demonstrated in model systems of alchemical changes in the Valine dipeptide and mutation cycle of the Alanine ↔ Valine ↔ Isoleucine (AVI) residues. Simulations showed that when GaMD boost potentials followed near-Gaussian distribution, the free energy change could be reweighted accurately through generalized cumulant expansion to the second order. The total free energy change often exhibited faster convergence using Selective GaMD (SGaMD) than using conventional MD (cMD). Accuracy of the free energy estimates from SGaMD-TI simulations was similar to or higher than those from cMD-TI simulations, although the differences were subtle for these small model systems. Meanwhile, dihedral angles in the model systems underwent significantly more frequent conformational transitions in SGaMD than in cMD, indicating improved sampling. Future studies are planned on larger systems with more complicated alchemical changes, such as ligand binding to proteins/nucleic acids and mutations at biomolecular binding interfaces. GaMD-TI should be broadly applicable to alchemical free energy calculations and therapeutic design.

**Significance Statement:** It is valuable to calculate alchemical free energy changes in drug design. We have integrated Gaussian accelerated Molecular Dynamics (GaMD) and Thermodynamics Integration (TI) for more efficient free energy calculations. Results that when GaMD boost potentials followed near-Gaussian distribution, the free energy change were reweighted accurately through generalized cumulant expansion to the second order. The total free energy change often exhibited faster convergence using Selective GaMD than using conventional MD, with significantly enhanced conformational sampling. Future studies are planned on larger systems with more complicated alchemical changes. GaMD-TI should be broadly applicable to alchemical free energy calculations.

## Introduction

Accurate prediction of free energies of ligand binding or alchemical changes remains a central challenge in computational chemistry/biophysics and drug discovery^1–3^. Reliable free energy calculations can substantially accelerate lead optimization, improve molecular design, and reduce experimental costs by providing quantitative estimates of binding affinities prior to synthesis and testing. Over the past several decades, rigorous statistical mechanics–based approaches, including alchemical free energy methods, have become indispensable tools for estimating relative and absolute binding free energies in biomolecular systems^2–5^. Among these approaches, Thermodynamic Integration (TI) has been established as one of the most widely used methods due to its strong theoretical foundation and broad applicability to ligand optimization and biomolecular transformations^6, 7^.

TI computes free energy differences by gradually transforming one molecular state into another through a series of intermediate alchemical states defined by a coupling parameter, λ^6, 7^. The free energy change is obtained by integrating ensemble averages of the derivative of the system Hamiltonian with respect to λ along the transformation pathway. When sufficient sampling is achieved at each λ state, TI can provide highly accurate free energy estimates. However, practical applications are often limited by the presence of large energy barriers, slow conformational transitions, and inadequate sampling of relevant configurational states in short simulations of each λ window. These challenges become particularly pronounced for flexible ligands, pocket residue flips, large biomolecular complexes, and systems exhibiting long-timescale conformational dynamics, leading to convergence difficulties and reduced accuracy.

A diverse variety of previous efforts have been designed to enhance the sampling and therefore improve the convergence and accuracy of TI and related forms of alchemical free energy calculations^8^. These approaches can broadly be described as methods using λ as a continuous variable^9–16^, methods using collective variable (CV)-based enhanced sampling algorithms such as metadynamics^10, 15–19^, umbrella sampling^9, 20, 21^, or adaptive biasing force^12, 15, 16, 22, 23^, and methods using CV-free enhanced sampling such as Hamiltonian replica exchange^8, 24–30^, expanded ensemble^14, 29, 31^, non-equilibrium switching^32–36^, and accelerated MD^30, 37^. These labels are not necessarily mutually exclusive – many methods describe λ as a continuous variable and use CV-based algorithms to bias the exploration of the alchemical coordinate^8–10, 12, 13, 15, 16, 23, 32^.

Gaussian accelerated Molecular Dynamics (GaMD) is an unconstrained enhanced sampling technique that adds a harmonic boost potential to smooth the system potential energy surface and facilitate barrier crossing^38–40^. GaMD enables efficient sampling of conformational space while preserving the ability to recover thermodynamic information through appropriate reweighting procedures. Previous studies have demonstrated the effectiveness of GaMD in accelerating biomolecular simulations and characterizing protein conformational changes, ligand binding mechanisms, and free energy landscapes across a wide range of biological systems^41^.

Inspired by previous studies combining the strengths of TI and enhanced sampling methods including accelerated MD^42^ and replica exchange^43, 44^, as well as results indicating that end state sampling via GaMD can alleviate common sources of error in TI^45^, we aim to integrate TI and GaMD as an attractive strategy for improving the accuracy and efficiency of free energy calculations. In this combined framework, GaMD enhances configurational sampling at each alchemical intermediate, reducing sampling errors associated with insufficient exploration of conformational space, while TI retains its rigorous statistical-mechanical formulation for calculating free energy differences. By improving convergence and reducing hysteresis along alchemical pathways, the GaMD-TI approach has the potential to deliver more reliable free energy predictions for complex biomolecular systems.

In this work, we present GaMD-TI, a hybrid methodology that combines enhanced conformational sampling from GaMD with rigorous alchemical free energy calculations using TI. The performance of GaMD-TI will be demonstrated on model system alchemical changes. The results show that GaMD-enhanced sampling improves convergence and accuracy of TI free energy estimates, highlighting the potential of GaMD-TI as a robust computational approach for free energy calculations in drug discovery and biomolecular design.

## Methods

### Gaussian accelerated Molecular Dynamics – Thermodynamic Integration (GaMD-TI) Theory

In Thermodynamic Integration (TI), the free energy change Δ*F* of an alchemical transformation can be calculated as:

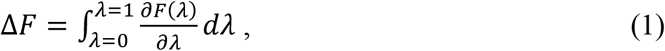

where λ is the alchemical parameter with λ = 0 and λ = 1 denoting the initial and final states, respectively (**Figure 1A**). Suppose the system potential *V* is a function of the coordinates of *N* atoms, *i.e.*, 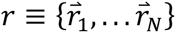. Then a derivative of the system free energy can be rewritten as:

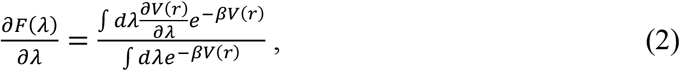

where β = 1/*k_B_T*, with *k_B_* as the Boltzmann constant and *T* as the temperature.

**Figure 1.**
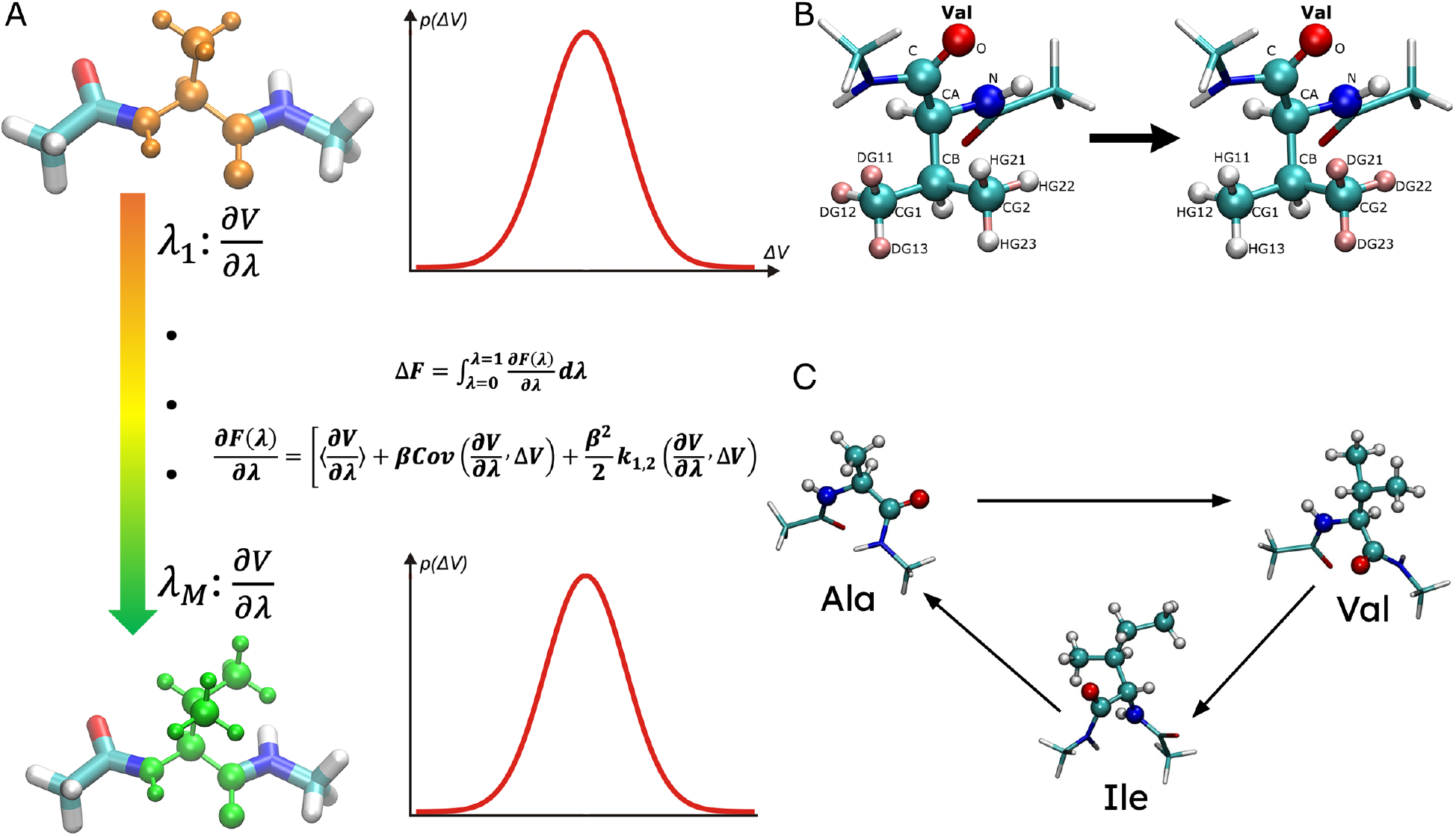
Schematic overview and model systems of GaMD-TI. (A) Alchemical change of a system from Initial State A (orange) to Final State B (green) is achieved through a number of *λ* windows (*λ*_1_, …*λ_M_*). In each *λ* window, when the GaMD boost potentials that are applied to enhance conformational sampling exhibit near-Gaussian distribution, the free energy 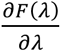 change can be approximated through generalized cumulant expansion to the second order. The total free energy change can be calculated by integrating 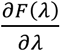 over the *λ* windows. (B) A model system of Val to Val (V2V) with H-D exchange the residue sidechain methyl groups. (C) The AVI mutation cycle system.

Gaussian accelerated Molecular Dynamics (GaMD) enhances the conformational sampling of biomolecules by adding a harmonic boost potential to reduce the system energy barriers. Details of the method have been described in previous studies^38, 46^. Here, we will apply a GaMD boost potential in each *λ* window to enhance the biomolecular conformational sampling when the system potential 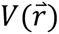 is lower than a threshold energy *E*:

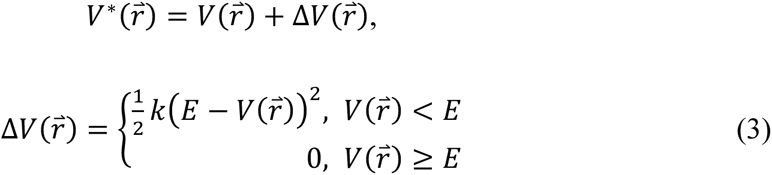

where *k* is the harmonic force constant. The two adjustable parameters *E* and *k* are automatically determined based on three enhanced sampling principles^38^. The threshold energy *E* needs to be set in the following range:

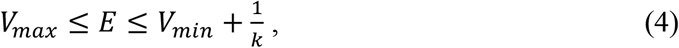

where *V_max_* and *V_min_* are the system minimum and maximum potential energies. To ensure that Eqn. (4) is valid, *k* must satisfy: 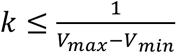. Let us define 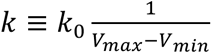, then 0 < *k*_0_ ≤ 1. The standard deviation of Δ*V* needs to be small enough (i.e., narrow distribution) to ensure proper energetic reweighting^47^: *σ*_Δ*V*_ = *k*(*E* − *V_avg_*)*σ_V_* ≤ *σ*_0_ where *V_avg_* and *σ_V_* are the average and standard deviation of the system potential energies, *σ*_Δ*V*_ is the standard deviation of Δ*V* with *σ*_0_ as a user-specified upper limit (e.g., 10*k_B_*T) for proper reweighting. When *E* is set to the lower bound *E=V_max_*, *k*_0_ can be calculated as:

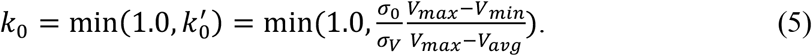

Alternatively, when the threshold energy *E* is set to its upper bound 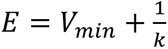 is set to:

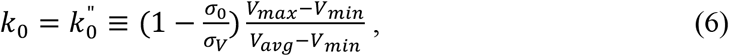

if 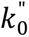 is found to be between *0* and *1*. Otherwise, *k*_0_ is calculated using Eqn. (5).

In GaMD-TI simulations with *V*(*r*) = *V*∗(*r*) − Δ*V*(*r*), we can rewrite Eqn. (2) as:

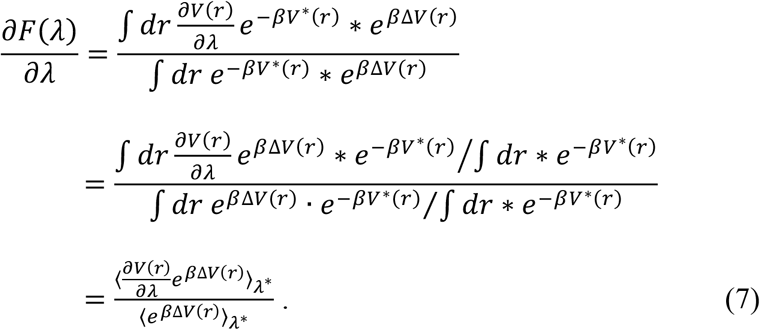

Next, we can approximate 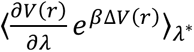, the ensemble average of 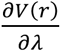 in a GaMD-boosted λ window, with generalized cumulant expansion. For simplicity, we assume 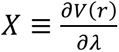 and 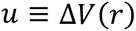. Then the generalized cumulant expansion can be applied as:

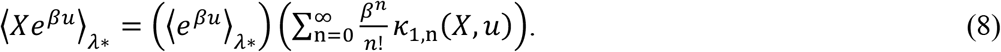

In the above expression, *κ*_1,n_(*X*, *u*) is the joint cumulant of order (1, *n*) between *X* and *u*. The first few terms are calculated as:

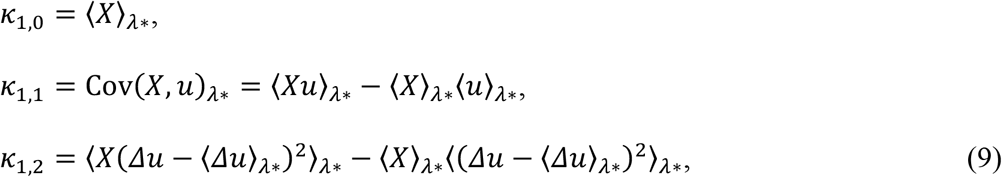

where *κ*_1,0_ is the average of *X*, *κ*_1,1_ is the covariance between *X* and *u*, measuring how they fluctuate together, and *κ*_1,2_ is a second-order correlation function. As the Gaussian boost potential usually exhibits a near-Gaussian distribution, cumulants higher than the 2nd order can be approximated to zero. Substituting these into the expansion gives:

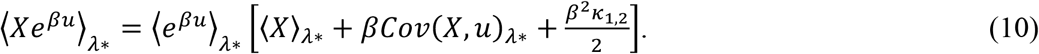

Then Eqn. (7) can be simplified as:

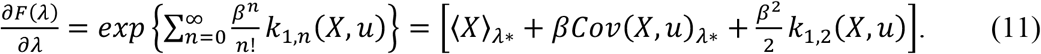

Therefore, we can calculate the total alchemical free energy by integrating the following 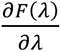 over the simulated λ windows (**Figure 1A**):

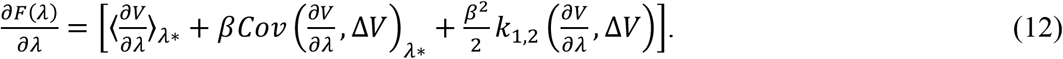

### Simulation Systems

In the Valine-to-Valine (V2V) model system (**Figure 1B**), three deuterium atoms (DG11, DG12, and DG13) bonded to the CG1 atom were alchemically changed to hydrogen, while three hydrogen atoms (HG21, HG22, and HG23) on the CG2 atom were changed to deuterium. Upon rotation of the residue sidechain dihedral angle χ_1_ (N–CA–CB–CG1), the CG1 and CG2 atom groups are equivalent. Provided sufficient conformational sampling of the Valine dipeptide, especially in the χ_1_ sidechain dihedral, the free energy change of the H–D exchange is in principle zero. We will use this as our first model system to test GaMD-TI.

The simulation structure of the V2V model system was prepared using the *tleap* module in AmberTools^48^. Two copies of a Valine dipeptide and a pre-equilibrated water box were loaded and combined into a single complex system. The Amber *ff14SB* force field^49^ was used for the dipeptide and the TIP3P model^50^ for water. A periodic box was assigned based on the coordinate centers of the combined unit. For TI, the combined topology and coordinates were processed using *tiMerge*^48^. The first and second copies of the Valine dipeptide were designated as the two end-state molecular regions, respectively. The Val residues with different side chains were specified as the corresponding TI regions that were described using soft-core (SC) potential functions for the alchemical transformation (**Figure 1B**). The merged dual-topology system was saved for subsequent TI simulations.

Dual-topology structures for the Alanine-to-Valine (A2V), Valine-to-Isoleucine (V2I), and Isoleucine-to-Alanine (I2A) mutation systems (**Figure 1C**) were prepared using the same process as with the V2V system using the *tleap* module and *tiMerge* in AmberTools^48^. The tLEaP *sequence* command was used to generate the residues with ACE and NME capping groups at the N- and C-termini, respectively. The Amber ff14SB and TIP3P force fields were used for the dipeptides and water, respectively.

### Simulation Protocols

For the V2V model system with the alchemical λ value set to 0.5, the system structure was initially energy minimized for 5000 steps and gradually heated to 300K with the constant-temperature, constant-volume ensemble (NVT) using a Langevin thermostat with a collision frequency of 5 ps⁻¹ over 1.6 ns. Nonbonded interactions were truncated at 9 Å. A 2 fs timestep was used in the MD simulations. Positional restraints of 1.0 kcal mol⁻¹ Å⁻² were applied to peptide backbone C, N, and CA atoms. This is followed by 8ns equilibration at 300K using the isothermal– isobaric (NPT) ensemble with the backbone constraints. Then the system is further equilibrated for 10ns at 300K using the NPT ensemble without any atom constraints. Next, the system alchemical parameter λ was gradually decreased from 0.5 to 0.43738, 0.31608, 0.20634, 0.11505, 0.04794, and 0.00922, and increased from 0.5 to 0.56262, 0.68392, 0.79366, 0.88495, 0.95206, and 0.99078 in two directions. Therefore, the TI simulations were performed in 12 different λ windows. In each window, the system is equilibrated for 2ns at 300K using the NPT ensemble without any atom constraints given the new λ value.

In each λ window, using the fully equilibrated structure, TI simulations of the V2V system proceeded with three independent runs using conventional MD (cMD) for 10 ns or different GaMD algorithms. The GaMD algorithms included the “Total-boost GaMD (GaMD_Tot)”, “Dihedral-boost GaMD (GaMD_Dih)”, “Dual-boost GaMD (GaMD_Dual)”, “Selective GaMD (SGaMD)”, and “Dual-boost SGaMD (SGaMD_Dual)”, as summarized in **Table 1**. In GaMD_Tot, the total potential energy of the system other than the TI/SC regions is boosted. In GaMD_Dih, only the dihedral energetic term of the system other than the TI/SC regions is boosted. In GaMD_Dual, both the dihedral and remaining total potential energies of the system other than the TI/SC regions are boosted. In SGaMD, only the total internal potential energy of the two TI regions described by SC potential functions is boosted. In SGaMD_Dual, both the total internal potential energy of the two TI/SC regions and the total potential energy of non-TI/SC regions of the system are boosted. It is important to note that 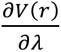 is not boosted in any of these GaMD-TI algorithms, which ensured proper reweighting of the simulations for free energy calculations. The GaMD-TI simulation of each λ window comprised of 0.4 ns short cMD to collect potential statistics, 1.6 ns GaMD equilibration after adding the boost potential(s), and then 10 ns GaMD production run. GaMD production frames were saved every 0.2 ps for analysis.

**Table 1.** The GaMD algorithms implemented in Amber26 for boosting TI simulations include the “Total-boost GaMD (GaMD_Tot)”, “Dihedral-boost GaMD (GaMD_Dih)”, “Dual-boost GaMD (GaMD_Dual)”, “Selective GaMD (SGaMD)”, and “Dual-boost SGaMD (SGaMD_Dual)”. In GaMD_Tot, the total potential energy of the system other than the TI/SC regions is boosted. In GaMD_Dih, only the dihedral energetic term of the system other than the TI/SC regions is boosted. In GaMD_Dual, both the dihedral and total potential energies of the system other than the TI/SC regions are boosted. In SGaMD, only the total internal potential energy of the two TI regions described by SC potential functions is boosted. In SGaMD_Dual, both the total internal potential energy of the two TI/SC regions and the total potential energy of non-TI/SC regions of the system are boosted. It is important to note that 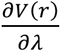 is not boosted in any of these GaMD-TI algorithms, which ensured proper reweighting of the simulations for free energy calculations.

| **GaMD Algorithm** | **Flag igamd** | **First Boosted Energy Term** | **Second Boosted Energy Term** |
| --- | --- | --- | --- |
| GaMD_Tot | 1 | pot_ene%total<br>[non-TI/SC] | None |
| GaMD_Dih | 2 | none | pot_ene%dihedral<br>[non-TI/SC] |
| GaMD_Dual | 3 | pot_ene%total<br>[non-TI/SC] | pot_ene%dihedral<br>[non-TI/SC] |
| SGaMD | 29 | ti_ene(1,si_pot_ene) +<br>ti_ene(2,si_pot_ene)<br>[SC] | None |
| SGaMD_Dual | 30 | ti_ene(1,si_pot_ene) +<br>ti_ene(2,si_pot_ene)<br>[SC] | pot_ene%total<br>[non-TI/SC] |

The A2V, V2I, and I2A systems were equilibrated using a forward and a reverse protocol which followed the procedure laid out in the Amber “Relaxation of Explicit Water Systems” tutorial. Each protocol was conducted on three independent replicates of the system. In the forward protocol, the alchemical λ value was set to 0.00922 and the system was minimized for 1000 cycles with full solute restraints with weight 100 kcal mol⁻¹ Å⁻², then heated from 100 K to 300 K for 1 ns with the same restraints, then 4 ns of constant pressure relaxation, followed by 1 ns relaxation with full solute restraints reduced to 10 kcal mol⁻¹ Å⁻². Then, 5000 cycles of minimization with backbone restraints at weight 10 kcal mol⁻¹ Å⁻², and a series of 1 ns constant pressure relaxation steps with backbone restraints reduced to 10, 1, and then 0.1 kcal mol⁻¹ Å⁻². Finally, 1 ns of unrestrained constant pressure relaxation and 4 ns of unrestrained constant volume relaxation. A 1 fs timestep was used for these relaxation steps and SHAKE was left off. In the reverse protocol, the same process was used except the λ value was set to 0.99078.

Next, in the forward protocol the equilibrated structure’s alchemical λ value was serially increased from 0.00922 to 0.99078 using the same 12-point Gaussian quadrature λ values mentioned for the V2V system. In the reverse protocol, the λ value was serially decreased from 0.99078 to 0.00922. In each window, the system was equilibrated for 1 ns at 300 K using NVT ensemble. Finally, the three replicates of the 12 λ-equilibrated structures were used as the starting points for 10 ns of cMD-TI as well as GaMD-TI. These simulations were run using the NVT ensemble with SHAKE on, a particle mesh Ewald nonbonded cutoff of 9 Å, a Langevin collision frequency of 1.0 ps⁻¹. In the GaMD-TI simulations, 2 ns of cMD-TI to collect boost potential statistics (ntcmd) and 2 ns of GaMD equilibration preceded the 10 ns of GaMD-TI production.

### Simulation Analysis

The VMD^51^ and CPPTRAJ^52^ tools were used for system visualization and simulation analysis. A Python script has been added in the *PyReweighting* package^38, 47, 53^ to reweight GaMD-TI simulations (termed “*PyReweighting-GaMD-TI*”: https://github.com/MiaoLab20/GaMD-TI) and calculate alchemical free energies. CPPTRAJ is used to calculate dihedral angles and examine conformational sampling of model systems in the simulation trajectories.

## Results

### GaMD-TI simulations show mostly narrow distributions and low anharmonicity of the boost potentials

TI simulations were performed on the V2V model system using cMD, Total-boost GaMD (GaMD_Tot), Dihedral-boost GaMD (GaMD_Dih), Dual-boost GaMD (GaMD_Dual), Selected GaMD (SGaMD) and Dual-boost SGaMD (SGaMD_Dual). The boost potential statistics and distribution anharmonicity were evaluated over the full alchemical transformation (**Figure 2**). GaMD_Tot and GaMD_Dual generated nearly λ-independent mean boost potentials of approximately 5–6 kcal mol⁻¹, whereas the dihedral boost in GaMD_Dih was negligible (**Figure 2A, 2C, and 2E**). The SGaMD and SGaMD_Dual protocols produced larger, more λ-dependent boosts; in particular, SGaMD displayed localized increases at selected λ windows (**Figure 2G and 2I**). The anharmonicity of the GaMD boost potentials remained low for GaMD_Tot, GaMD_Dual, and SGaMD_Dual (≈0.01–0.02, **Figure 2B, 2F and 2J**), while GaMD_Dih exhibited larger anharmonicity values (≈0.12–0.13, **Figure 2D**). SGaMD showed intermediate anharmonicity with a greater dependence on λ (**Figure 2H**).

**Figure 2.**
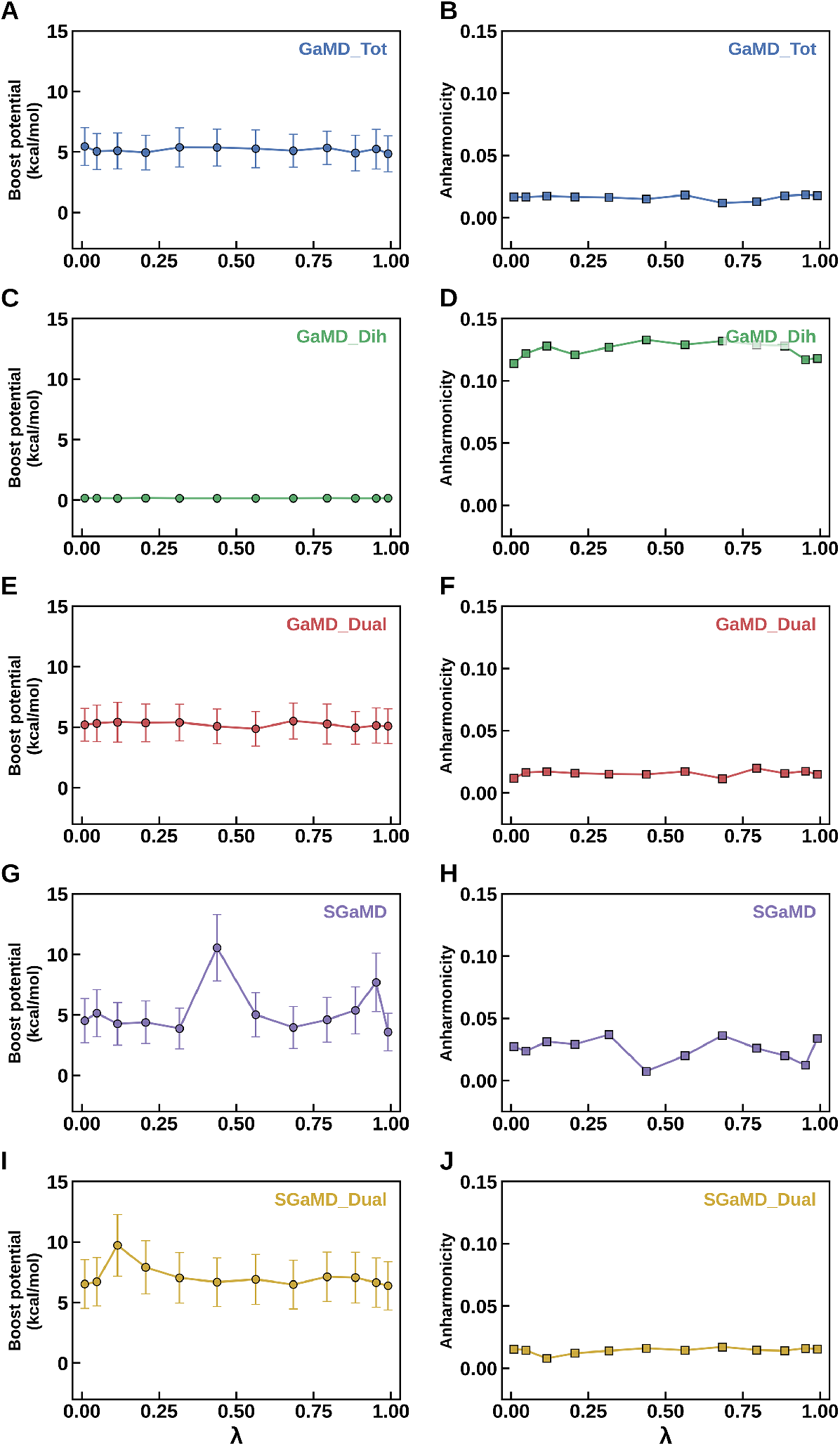
GaMD-TI simulations of the V2V model system show mostly narrow distributions and low anharmonicity of the boost potentials in each *λ* window, except for using GaMD_Dih. The average with standard deviations and anharmonicity of GaMD boost potentials calculated from a representative simulation using (A-B) GaMD_Tot, (C-D) GaMD_Dih, (E-F) GaMD_Dual, (G-H) SGaMD and (I-J) SGaMD_Dual.

The boost-potential distributions obtained with SGaMD and SGaMD_Dual were unimodal and approximately Gaussian over the representative λ windows examined (**Figure S1**). Their centers varied with λ, with SGaMD_Dual generally producing somewhat larger boost potentials. Consistent with the functional form of the boost, *ΔV* decreased smoothly with increasing system potential, *V(r)*, for both protocols (**Figure S2**). Except for GaMD_Dih, the approximately Gaussian distributions of the boost potentials in each λ window of TI simulations using other GaMD algorithms support the suitability of generalized cumulant expansion to the second order (“Gaussian approximation”) for energetic reweighting of the 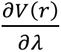 values (**Eqn. 12**). For GaMD_Dih, since the boost potentials are very small (**Figure 2C**), accurate energetic reweighting were achieved by calculating the “Exponential Average” reweighting factors directly (**Eqn. 7**).

### The free energy change in each *λ* window reweighted from GaMD-TI simulations agreed well with those of cMD-TI simulations

The TI integrands calculated from GaMD simulations with energetic reweighting closely reproduced those from cMD (**Figure 3**). For all methods including the GaMD_Tot, GaMD_Dih, and GaMD_Dual (**Figure 3A**), and the SGaMD and SGaMD_Dual (**Figure 3B**), ∂F(λ)/∂λ decreased from approximately 30 kcal mol⁻¹ at λ = 0 to negative values at intermediate λ, exhibited a small positive excursion near λ = 0.6–0.7, and decreased sharply to approximately −30 kcal mol⁻¹ at λ = 1. The close correspondence among the symmetric profiles of ∂F(λ)/∂λ vs. λ indicates that the GaMD enhanced-sampling protocols preserve the overall λ dependence of the TI integrand and the GaMD reweighted free energy values are highly accurate.

**Figure 3.**
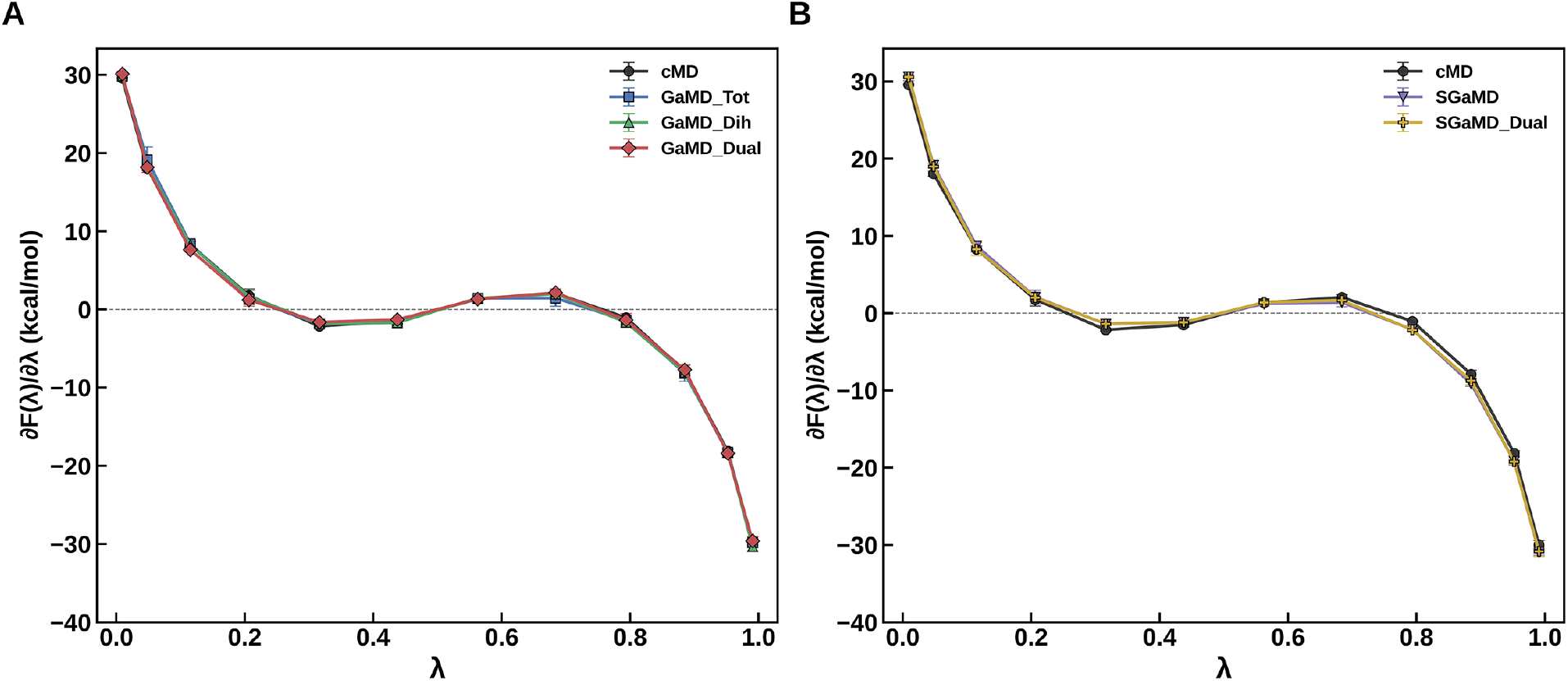
The free energy change in each *λ* window 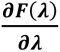 reweighted from GaMD-TI simulations agreed well with those of cMD-TI simulations on the V2V model system. The average 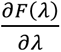 are plotted versus *λ* as calculated from three independent runs using (A) GaMD_Tot, GaMD_Dih and GaMD_Dual, and (B) SGaMD and SGaMD_Dual, with cMD as the reference.

### GaMD improved free energy convergence of the V2V model system

The V2V model system was constructed by exchanging the isotopic identities of the hydrogen and deuterium atoms on the two methyl groups of the valine side chain (**Figure 1B**). As expected for this symmetry-related transformation, the free energy change leveled off after several nanoseconds of simulation duration in the cMD-and GaMD-TI simulations over the 10-ns trajectories (**Figure 4B–F**). While the cMD-TI simulation approached to ∼0.1 kcal/mol after ∼4 ns (**Figure 4B**), the GaMD-TI simulations converged towards ∼0 kcal/mol as expected (**Figure 4B-F**). Notably, the SGaMD and SGaMD_Dual simulations exhibited faster convergence within ∼2-3 ns (**Figure 4E** and **4F**).

**Figure 4.**
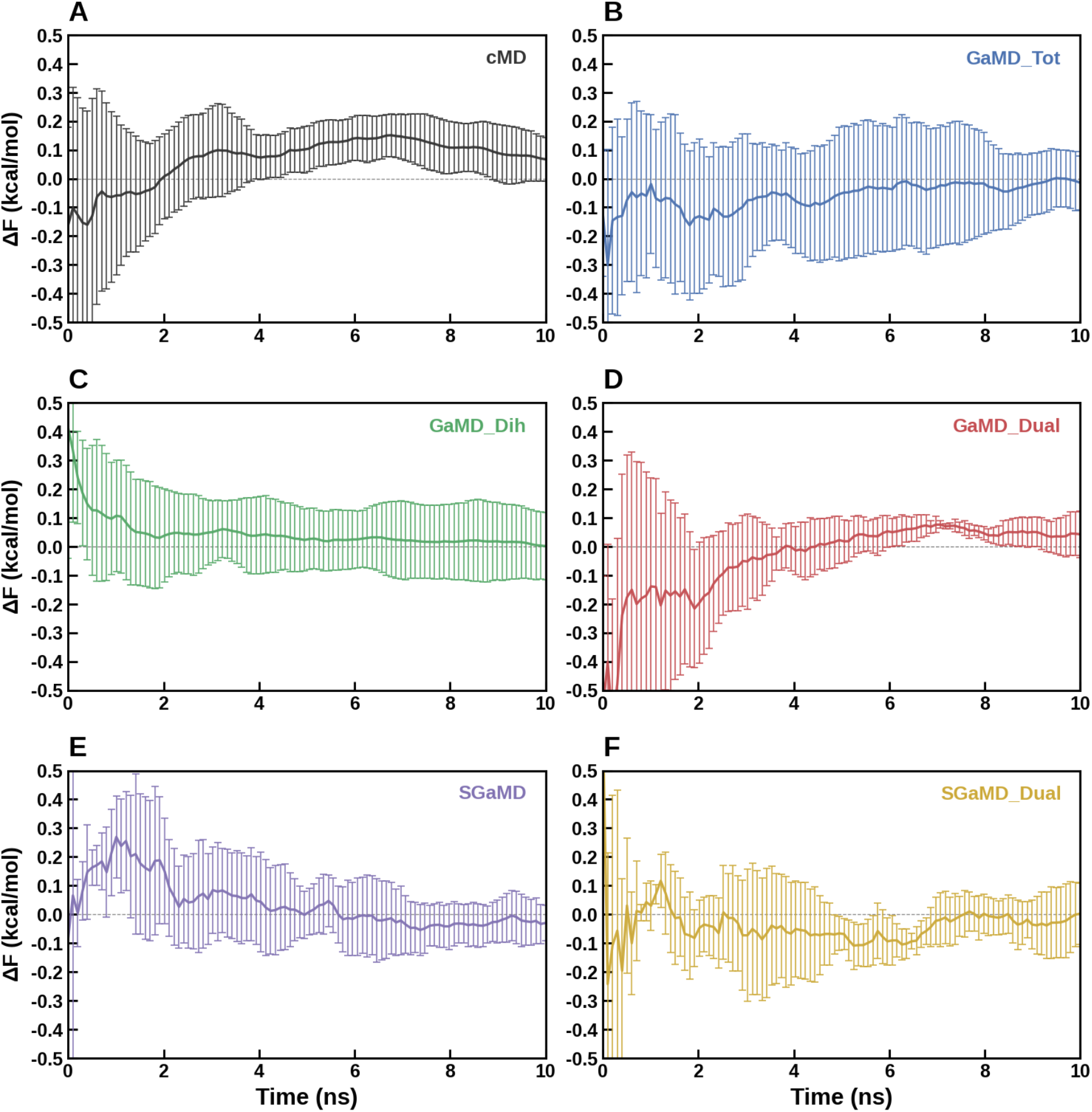
The total free energy change exhibits faster convergence and higher accuracy from GaMD-TI simulations than from cMD-TI simulations on the V2V model system. The total free energy change Δ*F* calculated from average of three independent simulations as a function of the simulation lengths using (A) cMD, (B) GaMD_Tot, (C) GaMD_Dih, (D) GaMD_Dual, (E) SGaMD and (F) SGaMD_Dual.

### SGaMD enhanced conformational sampling of the V2V model system

The χ₁ dihedral trajectories reveal marked differences in conformational sampling among the cMD-TI and GaMD-TI methods (**Figure 5**). In representative windows with *λ* = 0.01, 0.12, 0.32, and 0.56, cMD underwent relatively infrequent transitions between the rotameric basins (**Figure 5A**). In contrast, SGaMD and SGaMD_Dual exhibited repeated transitions among the principal rotameric states (**Figure 5B and 5C**). These results indicate that both SGaMD and SGaMD_Dual protocols significantly enhanced conformational sampling of the V2V model system (especially the side chain dihedral transitions) during the TI alchemical simulations, which contributed to faster convergence of the free energy calculations.

**Figure 5.**
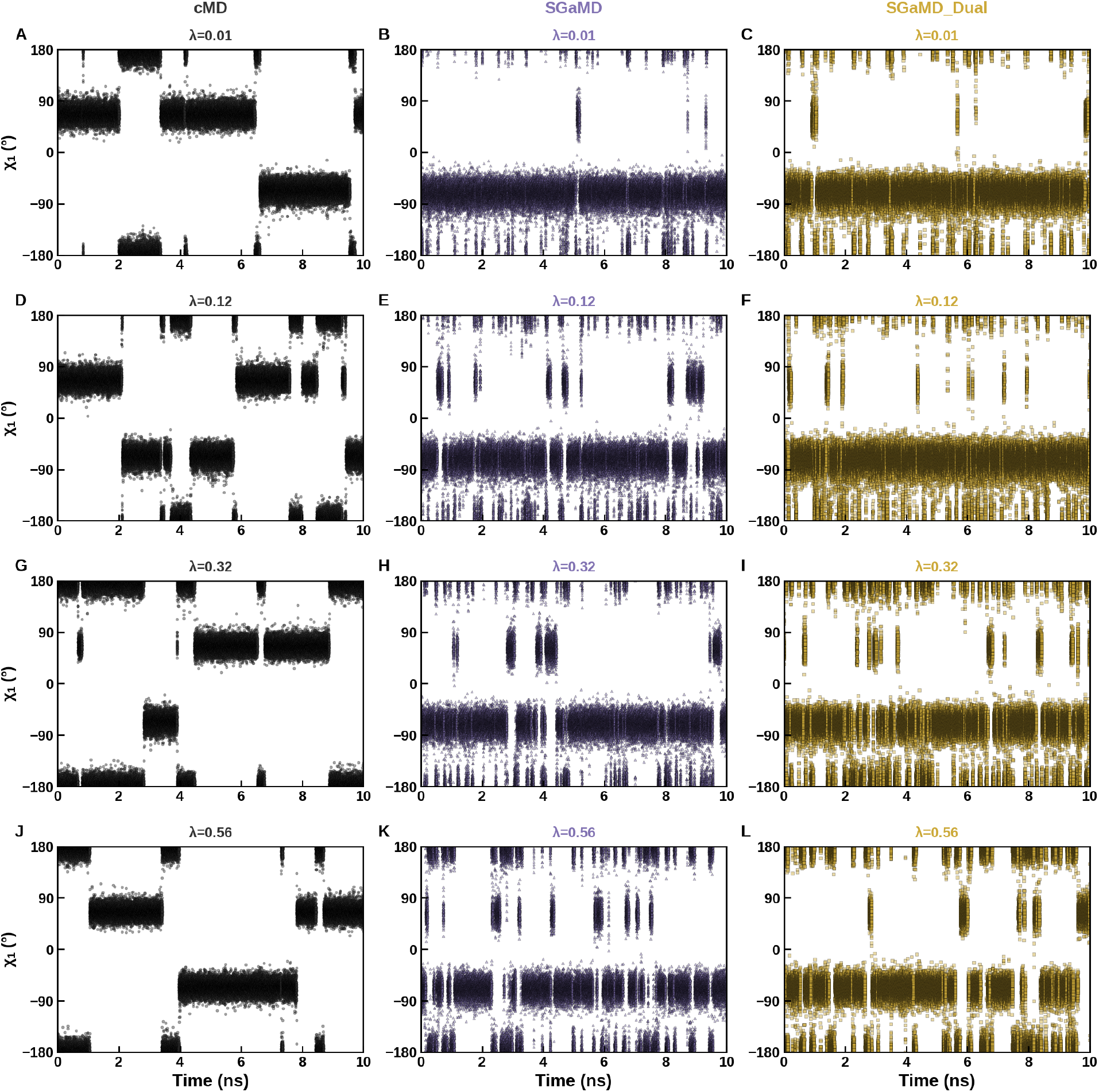
Dihedral conformational transitions were sampled more often in the SGaMD-TI simulations than from cMD-TI simulations on the V2V model system. The time courses dihedral angle χ_1_ calculated from a representative simulation using cMD, SGaMD and SGaMD_Dual in four example *λ* windows with (A-C) *λ* = 0.01, (D-F) *λ* = 0.12, (G-I) *λ* = 0.32, and (J-K) *λ* = 0.56.

### Free energy calculations of the AVI TI cycle

Next, we tested three non-zero free energy transformations with a cycle-closure that theoretically yields no free energy change: the A2V, V2I, and I2A model systems. For these experiments, we focused on SGaMD and SGaMD_Dual algorithms based on the performance observed in the previous V2V model system. As observed in the V2V model system, there was close correspondence in ∂F(λ)/∂λ for the individual 12 λ-windows between cMD-TI, SGaMD and SGaMD_Dual (**Figure 6**), suggesting that the GaMD reweighted free energy is highly accurate. It was gratifying then to see that the GaMD-TI algorithms faithfully recovered non-zero free energy estimates after reweighting with general cumulant expansion, and that despite the substantial boosts applied to the potential energy of these simulations, the results from both SGaMD and SGaMD_Dual were in good agreement with cMD-TI results (**Table 2**).

**Figure 6.**
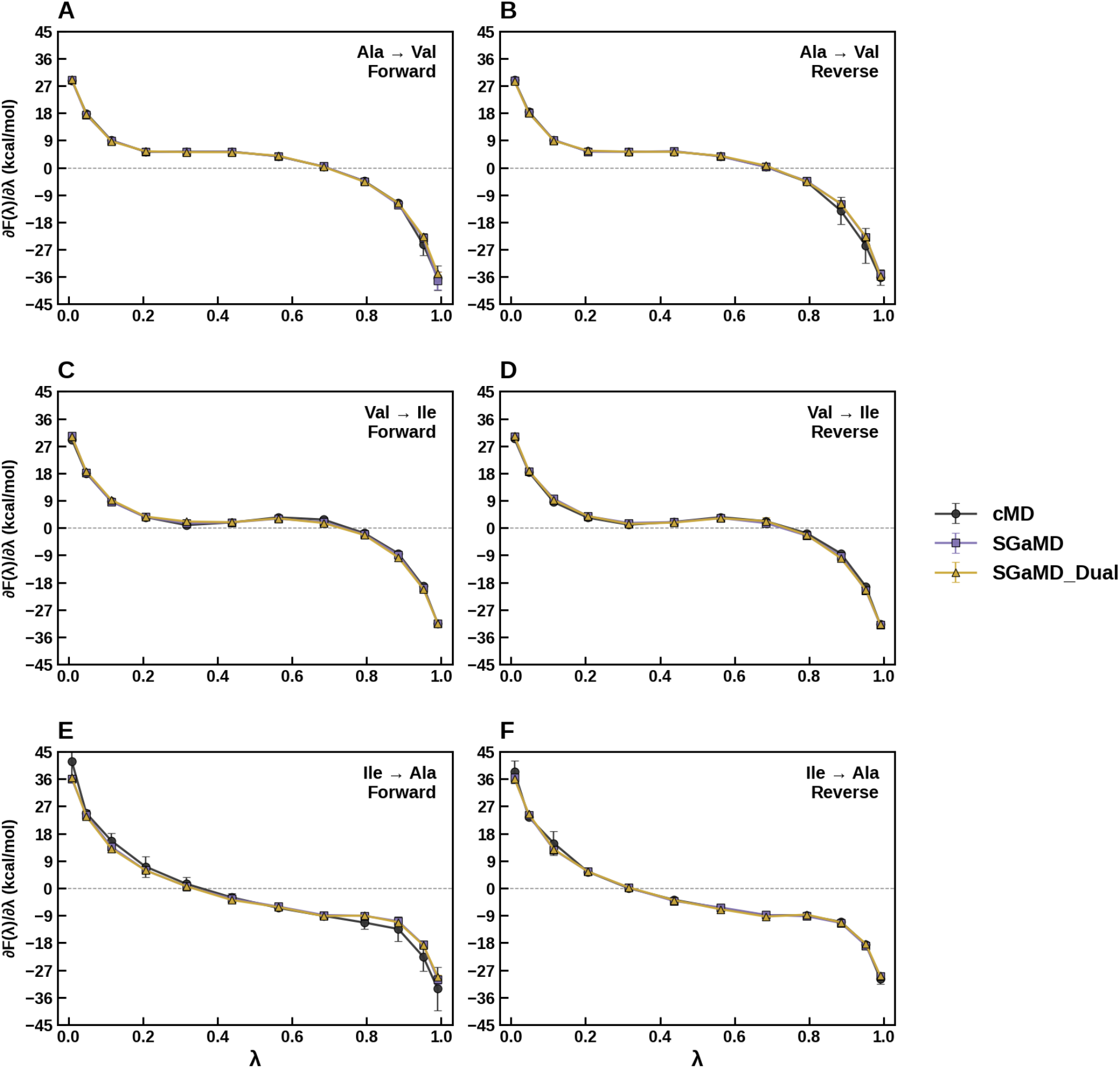
The mean derivative of potential energy with respect to λ (dF/dλ), reweighted by second cumulant expansion, for SGaMD (purple) and SGaMD_Dual (gold) agreed well with cMD-TI (black) for all λ windows in the AVI model system for both the forward (left column) and reverse (right column) transformations. First row is A2V, second row is V2I, third row is I2A. The mean was taken across three replicates, and the uncertainty in the error bars is the sample standard deviation of dF/dλ across the three replicates.

**Table 2.** Free energy changes for all test cases in the AVI model system. For each transformation the ΔF was computed by 12-point Gaussian quadrature integration and taking the average of three independent replicates. The uncertainties listed in the table are the standard error of the mean over the three replicates.

| Method | Mutation | $\Delta F$ forward (kcal/mol) | $\Delta F$ reverse (kcal/mol) | Hysteresis (kcal/mol) |
| --- | --- | --- | --- | --- |
| cMD-TI | A2V | $1.06 \pm 0.14$ | $0.87 \pm 0.36$ | $0.19 \pm 0.39$ |
| | V2I | $1.18 \pm 0.13$ | $1.14 \pm 0.14$ | $0.04 \pm 0.19$ |
| | I2A | $-2.02 \pm 0.17$ | $-1.95 \pm 0.32$ | $-0.07 \pm 0.36$ |
| **Cycle closure** |  | **$0.22 \pm 0.26$** | **$0.05 \pm 0.50$** |  |
| SGaMD | A2V | $1.08 \pm 0.03$ | $1.20 \pm 0.10$ | $-0.12 \pm 0.10$ |
| | V2I | $1.02 \pm 0.04$ | $1.02 \pm 0.07$ | $0.00 \pm 0.08$ |
| | I2A | $-1.82 \pm 0.09$ | $-2.16 \pm 0.19$ | $0.34 \pm 0.21$ |
| **Cycle closure** |  | **$0.28 \pm 0.10$** | **$0.06 \pm 0.23$** |  |
| SGaMD_Dual | A2V | $1.19 \pm 0.12$ | $1.25 \pm 0.04$ | $-0.06 \pm 0.13$ |
| | V2I | $1.05 \pm 0.06$ | $1.00 \pm 0.10$ | $0.05 \pm 0.12$ |
| | I2A | $-2.00 \pm 0.19$ | $-2.12 \pm 0.11$ | $0.12 \pm 0.22$ |
| **Cycle closure** |  | **$0.24 \pm 0.23$** | **$0.13 \pm 0.15$** |  |

To assist with evaluating convergence of the AVI simulations, we introduced mirrored TI simulations for each transformation (i.e. A2V, V2I, I2A) that used either the forward or reverse equilibrated structure as the starting point for the simulations. As shown in **Table 2**, the forward and reverse cycle-closures for GaMD-TI were more precise (i.e., tighter standard errors) with comparable accuracy when compared to cMD-TI. Forward cycle closure as a function of λ-window simulation time suggested rapid convergence for both GaMD-TI and cMD-TI, while reverse cycle closure painted a more nuanced picture with SGaMD-TI appearing to converge fastest, followed by cMD-TI and then SGaMD_Dual-TI (**Figure 7**). Like cycle-closure, the forward-backward hysteresis for GaMD-TI was more precise with comparable accuracy when compared to cMD-TI (**Figure 8**).

**Figure 7.**
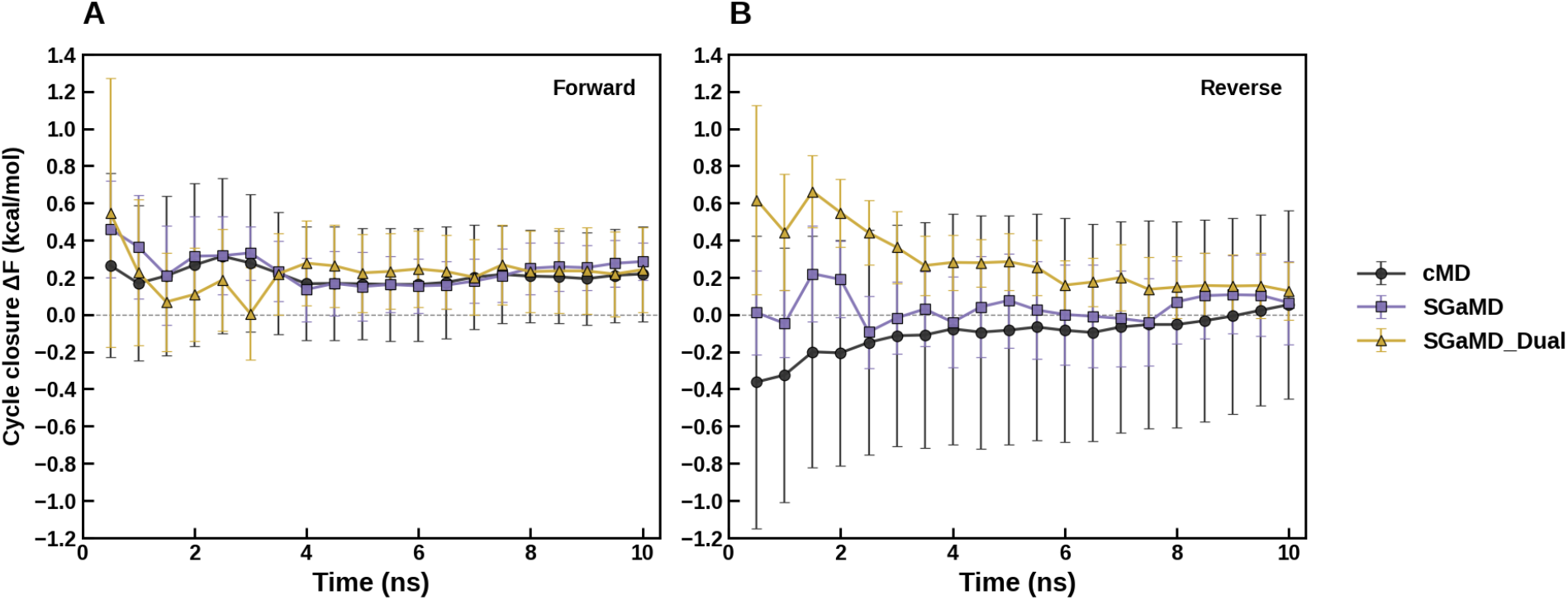
SGaMD and SGaMD_Dual exhibit comparable cycle closure accuracy and improved cycle closure precision when comprared to cMD-TI. The cycle closure, computed as the sum of the individual A2V, V2I, and I2A ΔF values along with the uncertainty (propagated standard error) is shown for the forward (left) and reverse (right) transformations as a function of simulation length for cMD-TI (black), SGaMD (purple), and SGaMD_Dual (gold).

**Figure 8.**
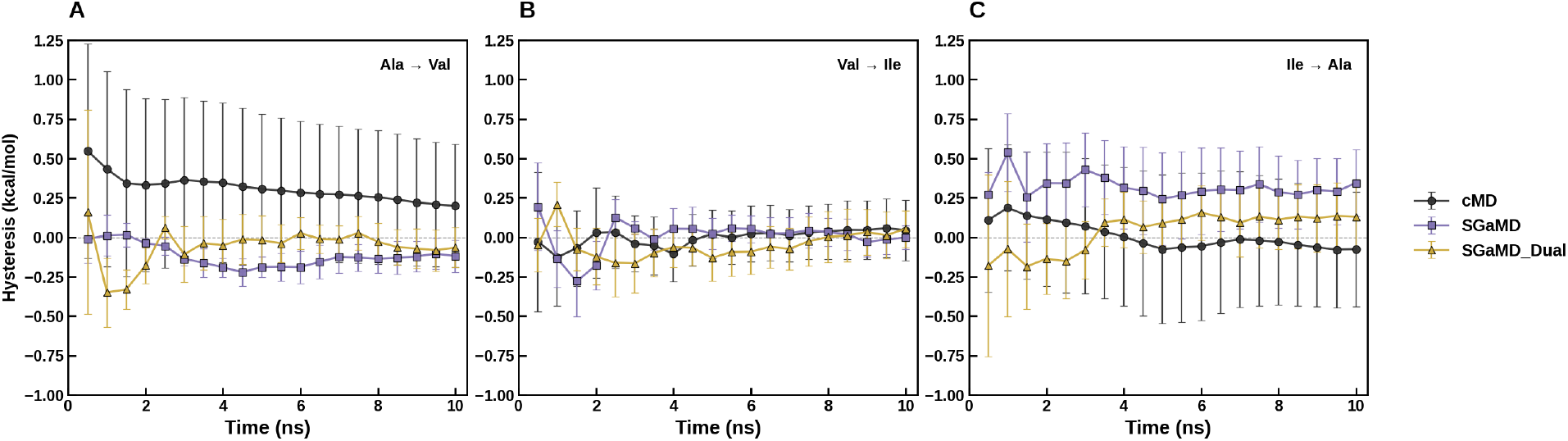
SGaMD and SGaMD_Dual demonstrate comparable hysteresis accuracy and improved hysteresis precision when compared to cMD-TI for the AVI system. The hysteresis, computed as the difference between the forward and reverse ΔF values along with the uncertainty as propagated standard error is shown as a function of simulation length for A2V (left), V2I (middle), and I2A (right) for cMD-TI (black), SGaMD (purple), and SGaMD_Dual (gold).

### Enhanced conformational sampling of the AVI model system

Analogous to the V2V system, the χ₁ dihedral trajectories in the AVI system exhibited marked differences between cMD-and GaMD-TI methods. **Figure 9** and **Figure 10** plot the Val and Ile χ₁ dihedral angles, respectively, for the V2I transformations with the different methods. While cMD eventually transitioned between the limited set of rotameric basins during a 10 ns simulation (**Figure 9 and 10**, left), increased transitions in corresponding GaMD-TI simulations were visually apparent for a majority of λ-windows as expected (**Figure 9 and 10**, middle and right), which likely contributed to the tighter precision observed with the GaMD-TI methods.

**Figure 9.**
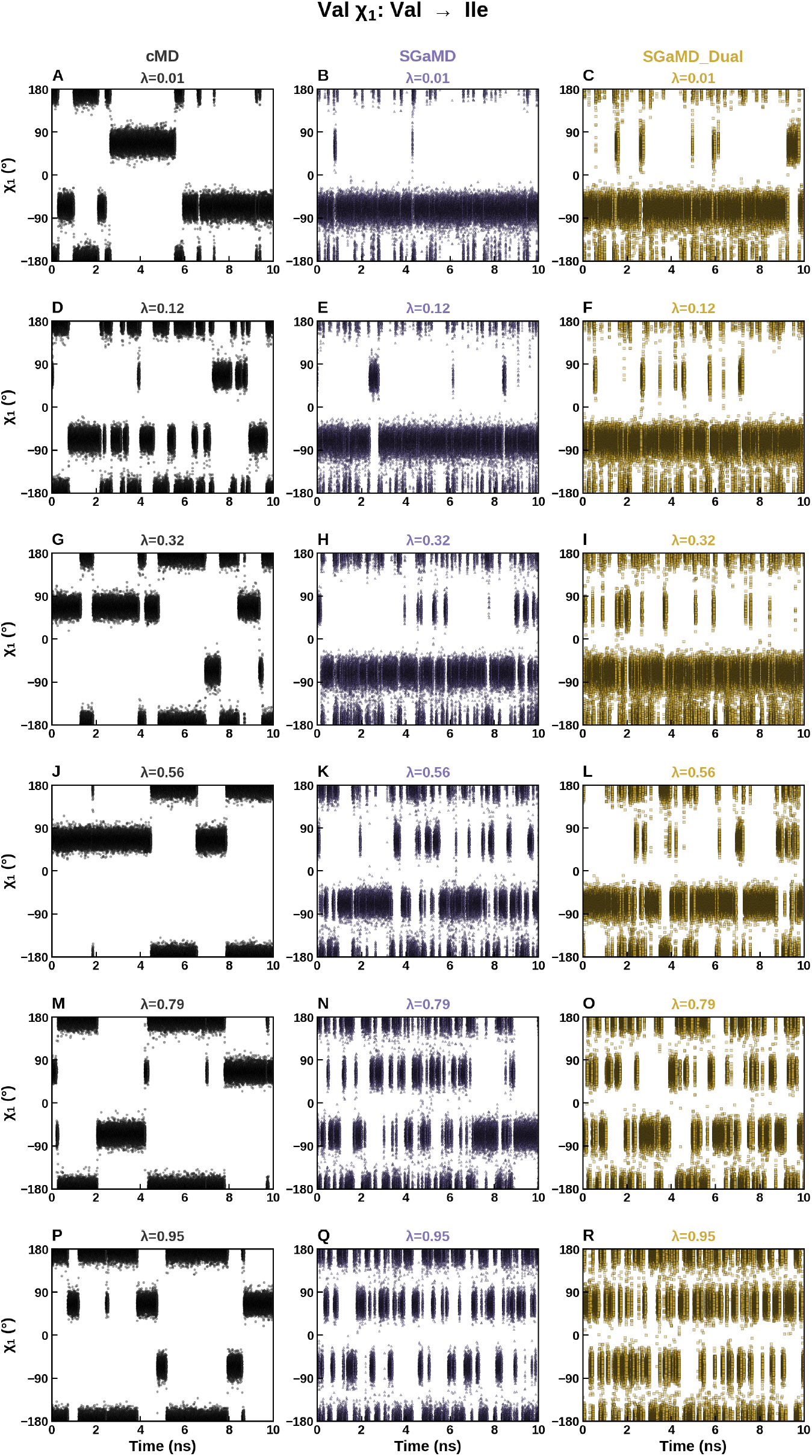
In the V2I system SGaMD and SGaMD_Dual enable enhanced sampling for the Val chi1 rotamer when compared to cMD-TI, except at λ values in which the Val topology is strongest (λ = 0.01, λ = 0.12), The Val chi1 angle (deg) over the course of the simulation is plotted for a representative replicate at representative λ values λ = 0.01 (first row), λ = 0.12 (second row), λ = 0.32 (third row), λ = 0.56 (fourth row), λ = 0.79 (fifth row), λ = 0.95 (sixth row) for cMD-TI (left column), SGaMD (middle column), and SGaMD_Dual (right column).

**Figure 10.**
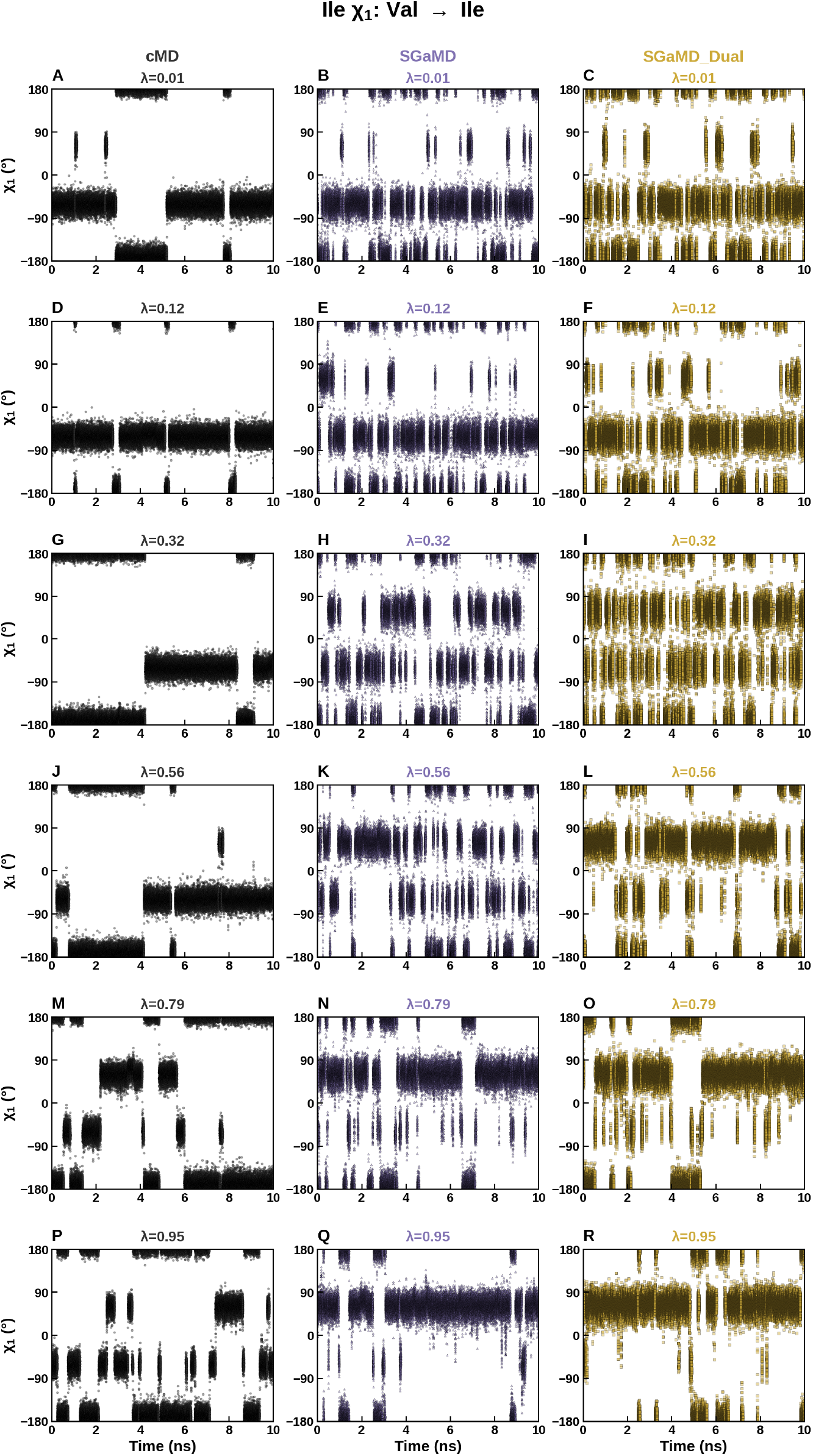
In the V2I system SGaMD and SGaMD_Dual enable enhanced sampling for the Ile chi1 rotamer when compared to cMD-TI, except at λ values in which the Ile topology is strongest (λ = 0.79, λ = 0.95). The Ile chi1 angle (deg) over the course of the simulation is plotted for a representative replicate at representative λ values λ = 0.01 (first row), λ = 0.12 (second row), λ = 0.32 (third row), λ = 0.56 (fourth row), λ = 0.79 (fifth row), λ = 0.95 (sixth row) for cMD-TI (left column), SGaMD (middle column), and SGaMD_Dual (right column).

## Discussion and Conclusions

In this study, we have successfully combined GaMD and TI to enhance conformational sampling and improve free energy calculations of alchemical changes. A novel formulation has been derived to accurately reweight the GaMD-TI simulations using generalized cumulant expansion to the second order when the GaMD boost potentials follow near-Gaussian distribution. This enables accurate reweighting of free energy changes in each λ window of TI simulations ∂F(λ)/∂λ, as well as the total free energy change Δ*F*.

Coupling TI with unconstrained enhanced sampling methods such as GaMD represents an attractive approach to improve the precision and accuracy of alchemical free energy predictions. Prior attempts to couple accelerated MD (aMD), a closely related predecessor to GaMD, and TI could be plagued by reweighting challenges that stemmed from exceedingly large boost potentials^37^. In this work, the reweighting problem was solved with two innovative augmentations: 1) by utilizing harmonic boosts characteristic of GaMD together with TI, which yielded near-Gaussian distributions of boost potential applied to dual topologies, and 2) by introducing a novel reweighting approach based on the generalized cumulant expansion to the second order.

GaMD-TI has been demonstrated on alchemical changes in the V2V and AVI model systems. Many more conformational transitions were observed for dihedral angles in the model systems in the SGaMD simulations than in cMD of the same lengths. This clearly indicates better enhanced sampling achieved using the SGaMD approach. Meanwhile, SGaMD-TI simulations could be accurately reweighted even with boosting potentials up to 10 kcal/mol. Moreover, increases in the boost potential magnitudes did not lead to larger free energy calculation errors as observed in SGaMD_Dual simulations. Overall, the total free energy change often exhibited faster convergence using SGaMD than cMD. Accuracy of the free energy estimates from SGaMD-TI simulations was also mostly similar to or higher than those from cMD-TI simulations.

Over the course of this study, five different types of GaMD-TI were explored. SGaMD and SGaMD_Dual yielded the most promising results, which may be attributable to their direct acceleration of the TI regions. The five GaMD-TI algorithms tested, however, were not intended to be exhaustive or comprehensive or final, and primarily serve to establish a proof-of-concept for GaMD-TI. In this initial iteration of GaMD-TI, we observed some curiosities that remain unaddressed and that may be idiosyncrasies of the particular type of GaMD-TI we chose to implement. For example, we noticed that while the majority of λ-windows clearly showed enhanced sampling of χ₁ dihedrals, SGaMD and SGaMD_Dual appeared less effective at accelerating conformational sampling when the topology containing the dihedral of interest was dominantly represented in the mixed potential (e.g., λ = 0.01 for Val χ₁ and λ = 0.95 for Ile χ₁ in V2I mutation system). This could be an artifact of the protocol which boosts based on the combined internal SC energies of TI region 1 and 2, potentially coupling two “non-interacting” parts of the system. Alternatively, this could be the result of exclusively boosting internal SC energy terms while omitting linearly scaled TI energy terms.

In the future, we plan to apply GaMD-TI simulations on larger systems with more complicated alchemical changes, such as ligand binding to proteins/nucleic acids and mutations at biomolecular binding interfaces. Moreover, further work could be carried out to optimize conformational sampling while continuing to preserve accurately reweighted free energies from GaMD-TI simulations. In this study, the GaMD boost being applied to either wholly the non-TI/non-SC region, or wholly within the TI/SC regions allowed for accurate computation of ∂F(λ)/∂λ without being disturbed by boost potential during the simulation. This also enabled accurate reweighting through generalized cumulant expansion to the second order since there was no correlation between the boost and ∂V(λ)/∂λ. However, this boosting strategy may not be enough to sample transitions between important dihedral angle degrees of freedom, such as those in the SC region which are constricted by high potential energy barriers due to their interactions with atoms outside of the SC region. It remains to be seen in future work how one can boost the interactions across the TI/SC boundary while still being able to recover accurate and reweighted ∂F(λ)/∂λ values.

Successful implementations of other unconstrained enhanced sampling methods and TI include Hamiltonian replica exchange between adjacent TI λ windows^24, 25, 27^ and ACES^28^. One key advantage of replica exchange is the ability to compute unbiased free energy quantities from the least perturbed replica exchange state. Algorithms bridging replica exchange with aMD demonstrated that this alleviated the significant challenge of energetic reweighting in aMD^30^. Although the SGaMD-TI and SGaMD_Dual-TI methods presented herein demonstrate the robust reweighting and recovery of accurate free energies, further integration with compatible techniques such as ACES and replica exchange could allow for even more powerful acceleration strategies, such as boosting interactions across the TI to non-TI boundary. GaMD-TI and related method developments will continue to improve free energy calculations.

## Supplementary Material

See the **Supplementary Material** for **Figures S1-S5** on the distributions of boost potentials Δ*V* and boost potentials Δ*V* vs. system potential *V(r)* calculated from GaMD-TI simulations of the V2V model system, as well as boost potentials applied in GaMD-TI simulations of the A2V, V2I, I2A model systems.

## Data Availability

The code used for building simulation systems, setting up simulations, running the simulations and analyzing the simulations of GaMD-TI have been provided through GitHub:

https://github.com/MiaoLab20/GaMD-TI.

GaMD-TI simulation files of the V2V model system can be downloaded at:

https://doi.org/10.6084/m9.figshare.33279687

https://doi.org/10.6084/m9.figshare.33283008

## Acknowledgements

This work used supercomputing resources with allocation award TG-MCB180049 through the Extreme Science and Engineering Discovery Environment ACCESS, which is supported by National Science Foundation grant number ACI-1548562, project M2874 through the National Energy Research Scientific Computing Center (NERSC), which is a U.S. Department of Energy Office of Science User Facility operated under Contract No. DE-AC02-05CH11231, and the UNC Research Computing. This work was also supported by Genentech (Project 6100098) and the startup funding project 27110 at the University of North Carolina-Chapel Hill.

## Declaration of the use of generative AI and AI-assisted technologies in manuscript preparation

The author(s) used ChatGPT in preparation of this manuscript. After using this tool/service, the author(s) reviewed and edited the content as needed and take(s) full responsibility for the content of the publication.

