## Supplementary Material for "Gaussian accelerated Molecular Dynamics – Thermodynamic Integration (GaMD-TI): Improved alchemical free energy calculations with enhanced sampling"

### Supplemental Material

**Fig. S1 Distributions of Boost potentials $\Delta V$ calculated from GaMD-TI simulations of the V2V model system in example $\lambda$ windows**. The distributions of GaMD boost potentials $\Delta V$ calculated from a representative simulation using GaMD_Tot, GaMD_Dih and GaMD_Dual with (A-C) $\lambda = 0.01$, (D-F) $\lambda = 0.12$, (G-I) $\lambda = 0.32$, and (J-L) $\lambda = 0.56$.

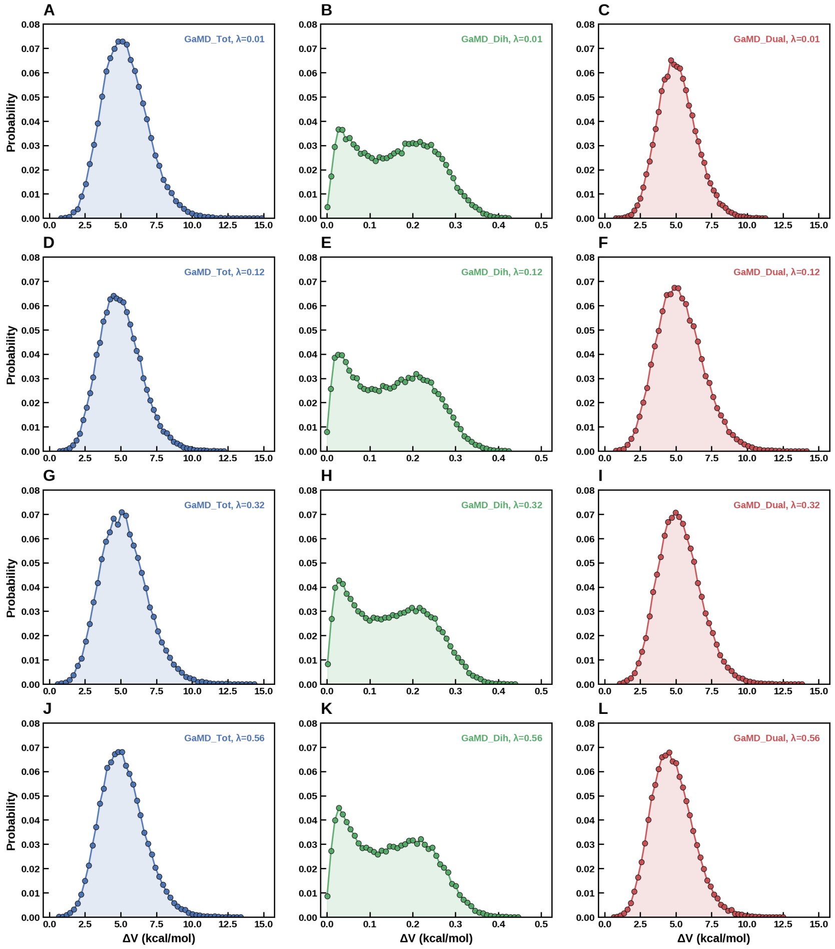

**Fig. S2 Distributions of boost potentials calculated from GaMD-TI simulations of the V2V model system exhibited near-Gaussian distributions in example $\lambda$ windows**. The distributions of GaMD boost potentials $\Delta V$ calculated from a representative simulation using SGaMD and SGaMD_Dual with (A-B) $\lambda = 0.01$, (C-D) $\lambda = 0.12$, (E-F) $\lambda = 0.32$, and (G-H) $\lambda = 0.56$.

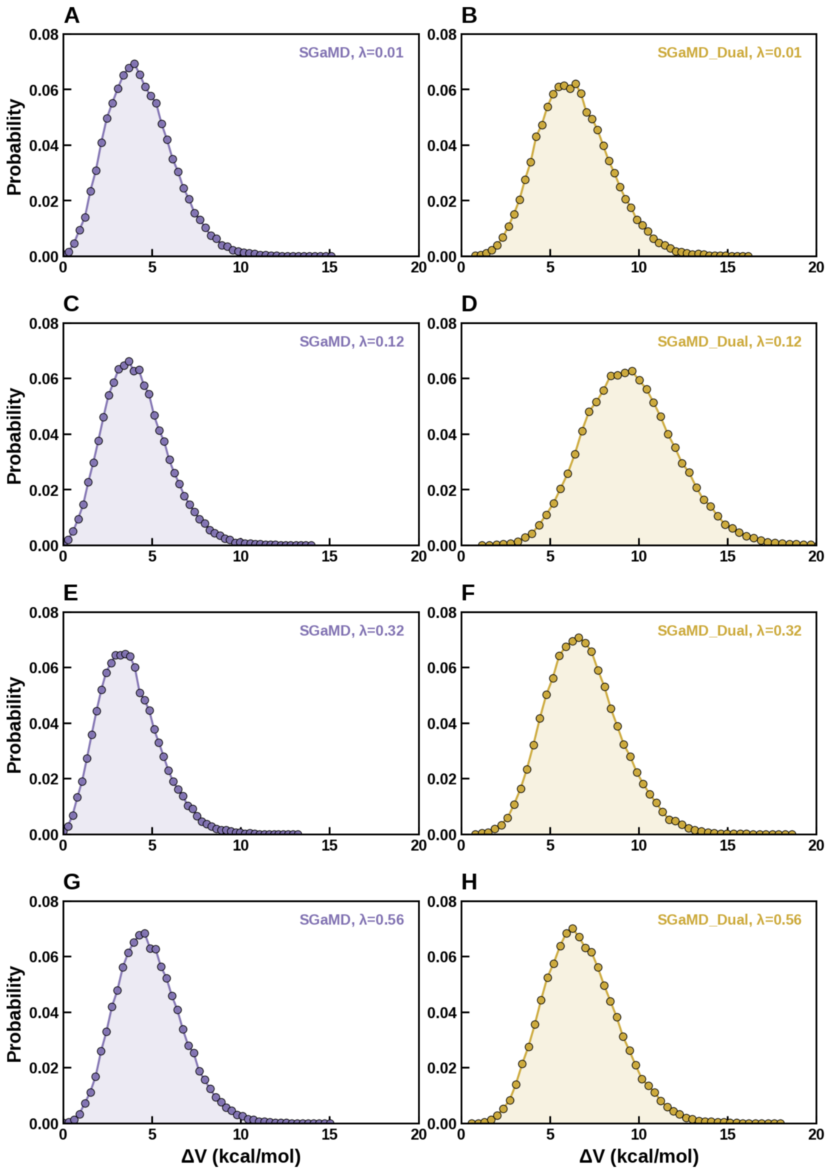

**Fig. S3 Boost potentials $\Delta V$ were applied higher with lower system potential *V(r)* in example $\lambda$ windows from GaMD-TI simulations on the V2V model system**. Boost potentials are plotted versus system potential from a representative simulation using SGaMD and SGaMD_Dual with (A-B) $\lambda = 0.01$, (C-D) $\lambda = 0.12$, (E-F) $\lambda = 0.32$, and (G-H) $\lambda = 0.56$.

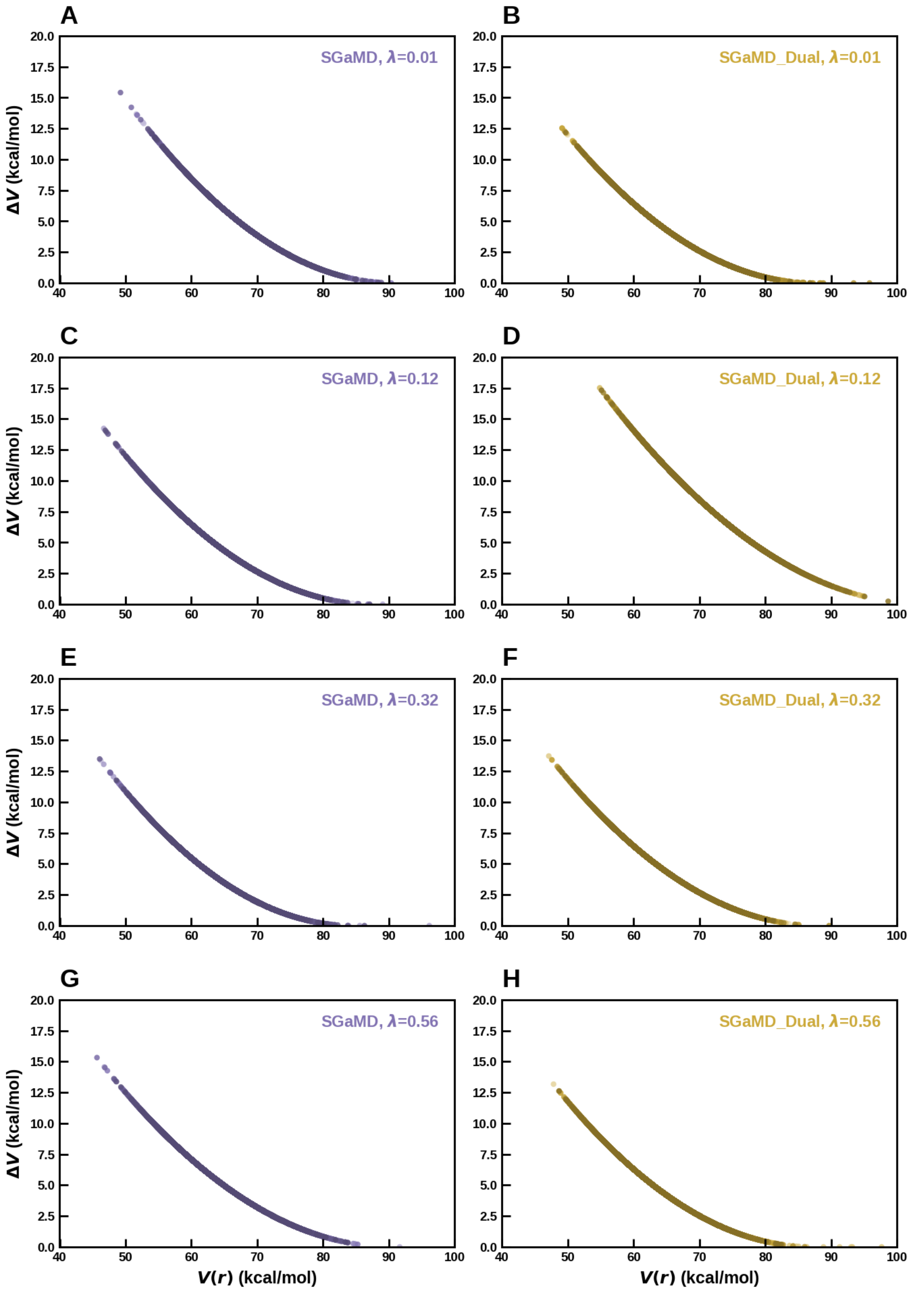

**Fig. S4 SGaMD adds approximately 4-6 kcal/mol of potential energy boost to the A2V, V2I, I2A model systems with sigma0P = 6.** Below, mean boost potential with SGaMD across three replicates with uncertainty (standard error of the mean) is plotted for A2V (first row), V2I (second row) and I2A (third row) for forward (left column) and reverse (right column) transformations

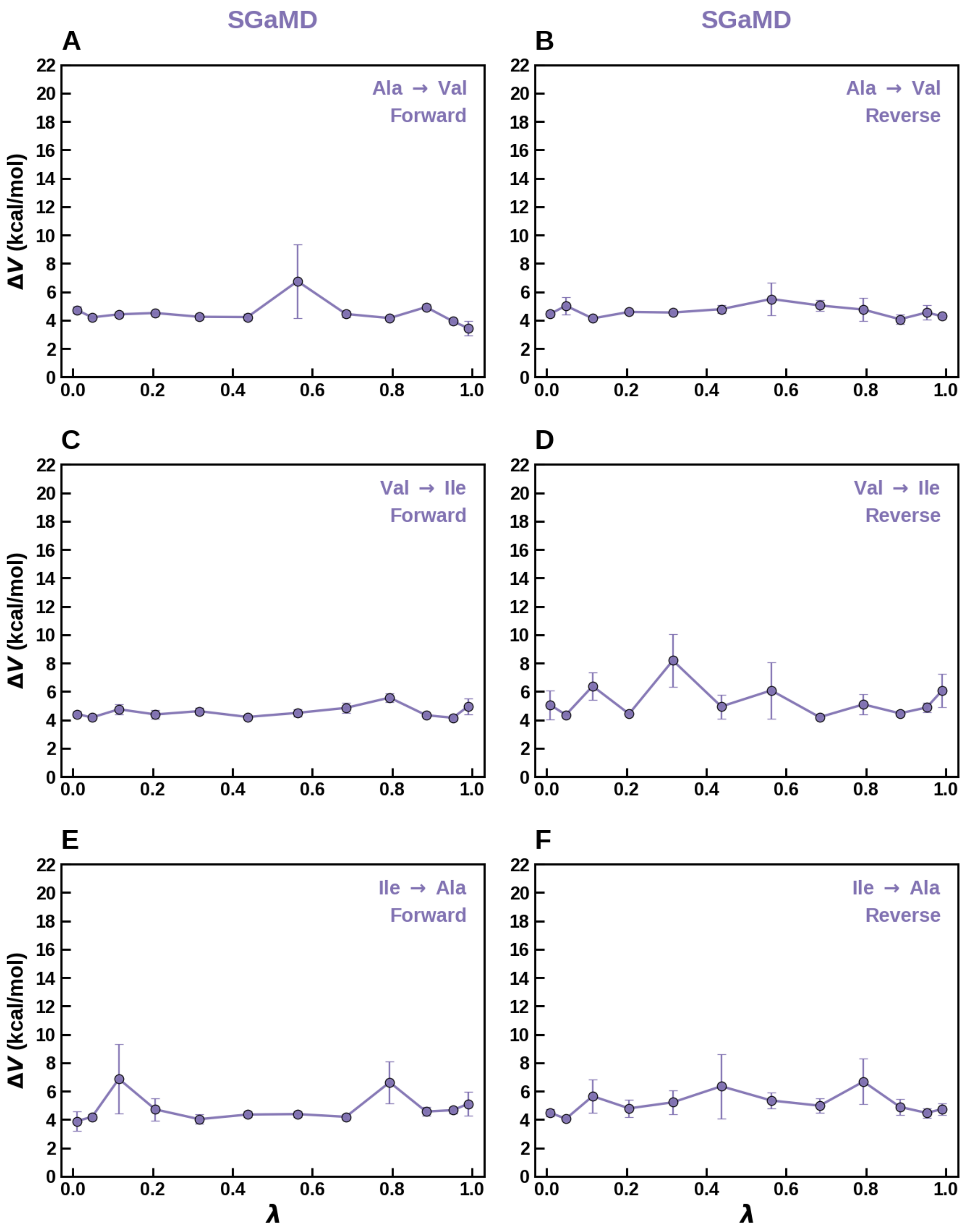

**Fig. S5 SGaMD_Dual adds approximately 5-10 kcal/mol of potential energy boost to the A2V, V2I, I2A model systems with sigma0P = 6, sigma0D = 1.** Below, mean boost potential with SGaMD_Dual across three replicates with uncertainty (standard error of the mean) is plotted for A2V (first row), V2I (second row) and I2A (third row) for forward (left column) and reverse (right column) transformations.

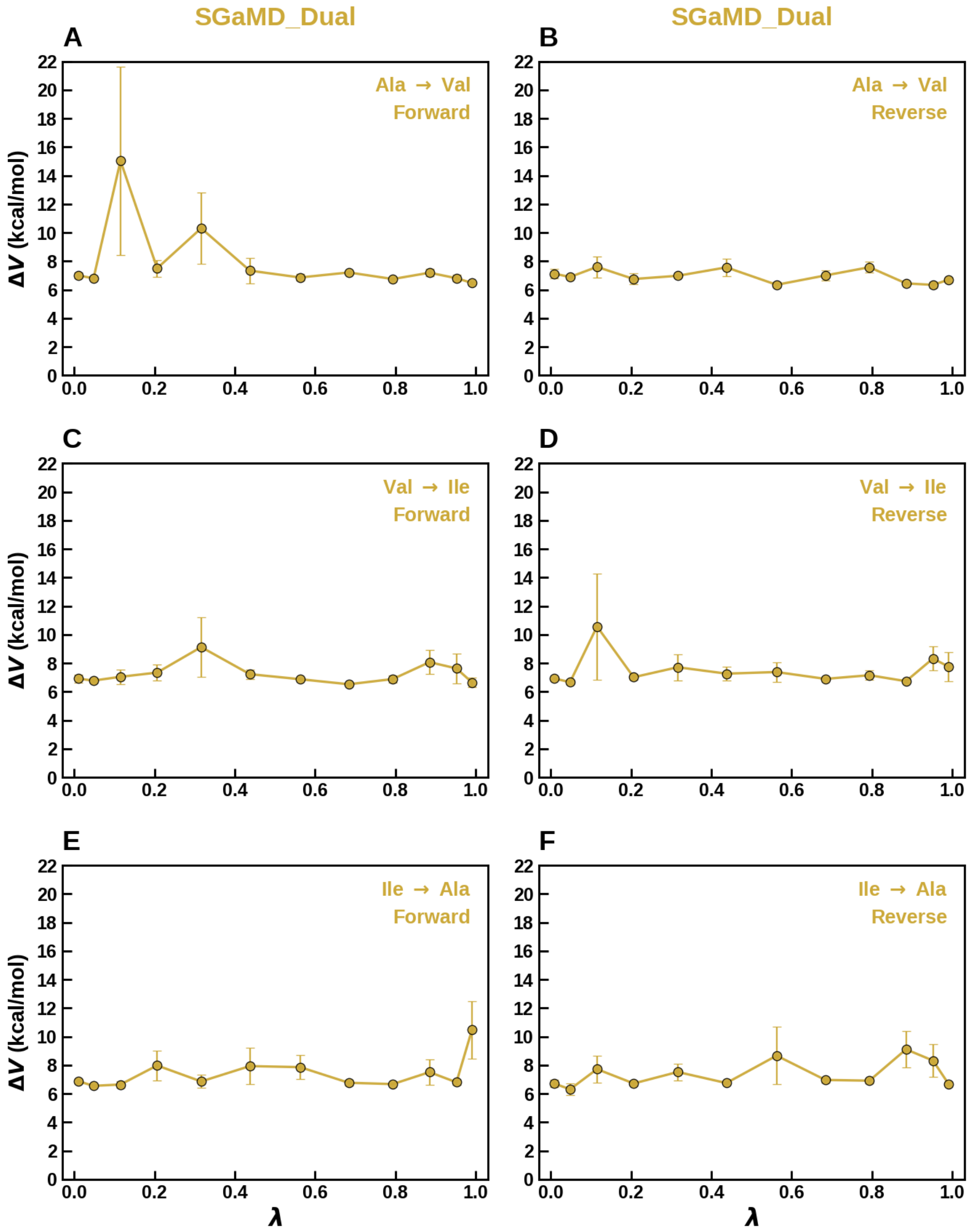
